# Genomic-inferred cross-selection metrics for multi-trait improvement in a recurrent selection breeding program

**DOI:** 10.1101/2023.10.29.564552

**Authors:** Sikiru Adeniyi Atanda, Nonoy Bandillo

**Affiliations:** Big Data Pipe Line Unit, North Dakota State University, Fargo, ND 58108-6050, USA; Department of Plant Sciences, North Dakota State University, Fargo, ND 58108-6050, USA

**Keywords:** Selection index, usefulness criterion, genomic prediction, genomic estimated breeding value, optimal haploid value, genetic gain, genetic drift, stochastic simulation, quantitative trait nucleotide, breeding cycle

## Abstract

The major drawback to the implementation of genomic selection in a breeding program is the reduction of additive genetic variance in the long term, primarily due to the Bulmer effect. Increasing genetic gain and retaining additive genetic variance requires optimizing the trade-off between the two competing factors. Our approach integrated index selection in the genomic infer cross-selection (GCS) methods. With this strategy, we identified optimal crosses that simultaneously maximize progeny performance and maintain genetic variance for multiple traits. Using a stochastic simulated recurrent breeding program over a 40-year period, we evaluated different GCS metrics with other factors, such as the number of parents, crosses, and progenies per cross, that influence genetic gain in a breeding program. Across all breeding scenarios, the posterior mean-variance consistently enhances genetic gain when compared to other metrics such as the usefulness criterion, optimal haploid value, mean genomic estimated breeding value, and mean index selection value of the superior parents. In addition, we provide a detailed strategy to optimize the number of parents, crosses, and progenies per cross that maximizes short- and long-term genetic gain in a breeding program.

## Introduction

Feeding the increasing world population requires doubling the current food production (van Dijk et al., 2021). Achieving this goal requires accelerating genetic gain within the constraints of a limited budget and resources. The traditional plant breeding scheme involves: i) parental selection for crossing to develop families; ii) creating homogeneous progenies within families via selfing or double haploid technology; iii) evaluating families in the nursery for morpho-agronomic and disease assessment; and iv) advancing superior genotypes through the yield testing stages (**Fig. 1**). In general, the scheme is considered long, taking several evaluation stages for breeding materials to be recycled as parents and varieties released as products (Santantonio et al. 2020). Reducing the breeding cycle time (the duration of time required to select parents back into the crossing block to create the next generation of families) has been identified as a key factor in accelerating genetic gain (Gaynor et al., 2017; Cobb et al., 2019; Santantonio & Robbins, 2020).

The advancement in genotyping technology, decreasing associated costs (Varshney et al., 2017; Atanda et al., 2021), as well as advances in statistical modeling and computing power, have spurred the widespread adoption of genomic selection (GS) (Bernardo & Yu, 2007; Meuwissen et al., 2001; Santantonio et al. 2020). GS utilizes DNA information to predict the genomic estimated breeding values (GEBV) of new genotypes. It has proven to be an innovative tool for expediting the breeding cycle time and reducing phenotyping expenses (Gorjanc et al., 2018; Beyene et al., 2019; Cobb et al., 2019; Santantonio et al. 2020; Atanda et al., 2021). This acceleration in genetic progress is attributed to its ability to identify superior parent genotypes for breeding at an earlier stage compared to traditional phenotypic methods (Gorjanc et al., 2018; Cobb et al., 2019; Atanda et al., 2021). However, the swift short-term genetic gain achieved through GS contributes to a faster reduction in genetic diversity in subsequent generations due to increased inbreeding (Jannink, 2010; Lin et al., 2016; Gaynor et al., 2017; Cobb et al., 2019; Santantonio & Robbins, 2020; Werner et al., 2023). The primary determinant of prediction accuracy in GS relies on the genetic relatedness between the training and the prediction set (Clark et al., 2012; Lee et al., 2017; Atanda et al., 2021). In other words, the superior genotypes selected through truncation selection are more likely to exhibit higher average coefficients of coancestry, resulting in higher inbreeding rates in each selection cycle. Several studies (Meuwissen, 1997; Mohammadi et al., 2015; Daetwyler et al., 2015; Akdemir & Sánchez, 2016; Lehermeier et al., 2017; Gorjanc et al., 2018; Allier et al., 2019), have suggested an alternative approach for sustainable genetic gain over both short and long terms in a breeding program. As opposed to interbreeding genotypes with the highest GEBVs (as in truncation selection), these strategies propose establishing crosses between genotypes based on a cross predicted usefulness or merit. Cross usefulness is a metric that optimizes the mean of the progenies and genetic variance within the bi-parental population (progenies that share the same parents from a single cross) (Schnell and Utz, 1975; Zhong & Jannink, 2007). For example, (Daetwyler et al., 2015) replaced the GEBV, which is the total sum of the additive marker effects, with the optimal haploid value (OHV) values that maximize haplotype complementarity from crossing parents. The authors reported an increase in long-term genetic gain in OHV-enabled parental combination selection compared to the mean GEBV of superior genotypes using stochastic simulation. However, OHV did not take into account linkage disequilibrium between quantitative trait loci (QTLs) and the challenge of optimal partitioning of the genome into predefined haplotype segments (Lehermeier et al., 2017; Müller et al., 2018). In another study, Lehermeier et al. (2017) proposed a novel deterministic approach to predict the additive progeny variance of a cross from the phenotypic and genotypic information of the parents. The predicted additive progeny variance was used within the statistical framework of the usefulness criterion (UC) proposed by Schnell and Utz (1975) to select parent combinations for crossing blocks. In general, these metrics and others are typically evaluated based on individual traits. However, in practice, potential parents often possess multiple traits of economic and agronomic significance (Cerón-Rojas & Crossa, 2022; Wellmann, 2023). These traits have attributes linked to productive performance, adaptability, and production stability. To improve multiple traits simultaneously, selection index methods are commonly employed. These methods combine all relevant traits into a single index and prove highly useful for improving multiple traits with the desired selection response (Hazel & Lush, 1942; Rocha et al., 2018; Céron-Rojas et al., 2018; Cerón-Rojas & Crossa, 2022).

The Smith and Hazel selection index, which integrates genetic correlation with economic weights, has gained wide traction in animal breeding (Bernardo, 2020; Cerón-Rojas & Crossa, 2022; Wellmann, 2023). Determining suitable weights for different agronomic and quality traits remains a significant challenge, limiting the widespread adoption of this index in plant breeding. In this study, we consider the non-parametric rank summation index proposed by Mulamba and Mock (1978). It offers the distinct advantage of not requiring economic weights to compute the index for different genotypes (Smiderle et al., 2019; Casagrande et al., 2022). The rank summation index is based on ranking genotypes in relation to the desired trait and summing up the ranks for multiple traits simultaneously (Mulamba & Mock 1978; Cruz, 2003; Coutinho et al., 2019; Casagrande et al., 2022). Theoretically, selection on this index (a hypothetical new phenotype) should result in simultaneous improvements across all desired traits. To our knowledge, this is the first time index selection will be utilized within the framework of GP to select genotypes and parental combinations for crossing blocks. While our objective aligns with (Wolfe et al. 2021), our approach to achieving this goal is notably distinct. In addition, this study aims to identify the optimal number of parents, crosses, and progenies per cross in the North Dakota State University (NDSU) pulse breeding program using stochastic genetic simulation in the R package AlphaSimR (Gaynor et al., 2021).

## Materials and Methods

### Founder population and genetic parameters

A pea (*Pisum sativum* L.) genome size (cM) and chromosome sizes described in (Kreplak et al. 2019) were simulated using the Markovian Coalescent Simulator (MaCS) (Chen et al. 2009) implemented in AlphasimR (Gaynor et al., 2021). This resulted in a founder population of 200 non-inbred individuals with 7 chromosome pairs each.

In the founding population, we assumed that 2,100 segregating sites were evenly distributed across the chromosomes. From these sites, 500 segregating sites were randomly sampled per chromosome to serve as quantitative trait nucleotides (QTN) and single-nucleotide polymorphisms (SNPs). We simulated four polygenic traits: grain yield (YLD), 1000 kernel weight (TKW), days to physiological maturity (DPM), and plant height (PH). In quantitative genetic theory, it is assumed that the number of segregating QTN for polygenic traits will exceed the number of independent chromosome segments (*M_e_*) (Daetwyler et al., 2010, Bernardo, 2020). Pea has limited genetic diversity (Yang et al., 2022) and presumably has *M_e_* less than the 500 random QTN selected in our study. This aligns with several simulation studies that predominantly assume polygenic traits are controlled by 500 or greater QTN (Wientjes et al., 2015; Yao et al., 2018; Peters et al., 2020; Li et al., 2022; Sabadin et al., 2022).

Each QTN was assigned an additive effect that was sampled from a Gaussian distribution with a mean and variance obtained from variance components estimated from a multivariate model fitted to NDSU historical field yield trials. The means were (YLD = 5.78, TKW = 433.00, DPM = 81.00, PH = 67.00), and the variances were (YLD = 3.59, TKW = 50.10, DPM = 12.24, PH = 15.80). For simplicity, similar to (Li et al., 2022) we also omitted dominance and epistasis effects in the simulation.

### Phenotypes Simulation

Random noise sampled from a normal distribution with a mean of 0 and the error variance for the traits were added to the genetic values of the founder lines to produce the phenotype. The error variance was varied to reflect the plot level heritability currently obtained in the breeding program for the yield testing stages. Entry-mean narrow-sense heritability was set to 0.1 in the nursery stage for visual selection similar to Gaynor et al. (2017). The simulated genetic correlation between traits (off-diagonal element) and broad-sense heritability (diagonal element) were presented in (**Supp. 1**). The genotype-by-environment variance provided a non-heritable variation attributed to the locations; for detailed implementation in AlphaSimR, see (Gaynor et al., 2021) for details.

### Simulation Parameters

The simulation was based on the NDSU pea breeding program with several simulated treatments (**Table 1**). In all treatment scenarios, the number of individuals in the F_2_ generation was restricted to 15,000, while in the progeny row or nursery, the limit was set at 4,000. We developed a grid to evaluate different numbers of parents: 30, 40, and 50, and 50, 100, 150, and 200 crosses, respectively. The number of progenies per cross was limited to 300, 150, 100, and 75, respectively. Thus, the number of F_2_ individuals (15,000), which is the number of crosses multiplied by the number of progenies per cross, is constant across treatments. We used the UC, posterior mean-variance (PMV), OHV, mean GEBV, and mean index selection of the superior parents (denoted as MeanPheno) as metrics to choose parent pairings for crossing. In total, 60 simulation treatments were examined.

**Table 1:** Summary of the combination of the of number of parents, crosses and progenies per cross used for the simulation study.

| Number of parents | Number of crosses | Progenies per cross |
| --- | --- | --- |
| **30** | 50 | 300 |
|  | 100 | 150 |
|  | 150 | 100 |
|  | 200 | 75 |
| **40** | 50 | 300 |
|  | 100 | 150 |
|  | 150 | 100 |
|  | 200 | 75 |
| **50** | 50 | 300 |
|  | 100 | 150 |
|  | 150 | 100 |
|  | 200 | 75 |

Note: Multiplying the number of unique combinations, which is 12, by the number of cross-selection metrics (UC, OHV, PMV, MeanGEBV and MeanPheno) yields a total of 60 treatments.

### Simulation scenario

We utilized the same founder population across all treatments or breeding scenarios. Identification of superior genotypes as parents was performed based on the rank summation index (Mulamba and Mock, 1978). The genotypes are transformed into ranks based on their genotypic values according to the interest of the breeding program in increasing or decreasing the mean value of the trait. After obtaining the ranks for each genotype, we calculated the index as follows:

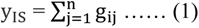

where y_IS_ is the index for genotype i and g_ij_ is the rank of genotype i for trait j.

The index served as a measure of the genotypes genetic value and was utilized as a derived phenotype. Using only 3,500 non-QTN markers, we fitted a whole-genome regression model (Eq. 2) using the derived phenotype (y_IS_) as a response variable to estimate the GEBV of the genotypes. Thus, superior genotypes were selected as parents based on the new phenotypic value and GEBV. We calculated the Rogers’ distance based on marker data between all possible combinations of the selected parents. Only parent combinations with a genetic distance less than 0.1 were preselected before the average value of the index selection (MeanPheno) and MeanGEBV were used as final decision metrics. This step was necessary to best simulate a typical procedure in a breeding program. For the UC, PMV, and OHV, there was no prior selection of crosses.

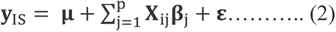

where: p is the marker size, **X**_ij_ is the genotype score at the j-th locus/QTN of the genome in the i-th line, 0 is the homozygous copies of the allele, 1 is the heterozygous copies of the allele, and 2 is the homozygous copies of the second allele, **β**_j_ is the effect of marker j-th on **y**_IS_. The markers effect was assumed to be independent and identical with Gaussian distributions; 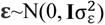. Also the residual error was assumed to be independent and identical with Gaussian distributions 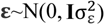. The whole-genome regression model was fitted with Bayesian Ridge Regression implemented in BGLR package (Pérez and de los Campos 2014). This assumed scaled inverse-*X*^2^ prior distributions assigned to the markers effect and residual variance (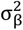 and 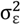) respectively. Samples from the posterior distribution were generated using the Markov chain Monte Carlo (MCMC) algorithm implemented in the BGLR package. We used 40,000 iterations, discarded the first 10,000 as burn-in and thinned to every 10th sample.

The estimated GEBV (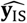) is the product of estimated markers effect 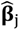 and allele dosages.

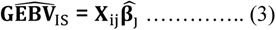

Mean GEBV was obtained as follows:

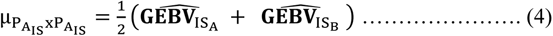

The formula for the expected variance of progeny for each P_A_ *X* P_B_ combination was formulated as proposed by Lehermeier et al. (2017):

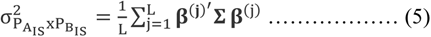

L is the size of the posterior sample postburn-in, **β**^(j)^ is the j-th thinned postburn-in sample of the MCMC algorithm from the whole-genome regression model (Eq. 2) and **Σ** is the variance covariance matrix between DH parents P_A_ *X* P_B_ alleles at QTN in progeny see Lehermeier et al. (2017) for details.

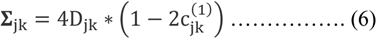

D_ij_ is the linkage disequilibrium (LD) parameter between alleles at loci j and k for parents P_A_ *X* P_B_. The parameter D_ij_ would be 0 if both parental pairs share the same allele at either locus j or k. Alternatively, it assumes a value of 0.25 or −0.25, depending on the linkage phase of the parental pair. c_jk_ is the recombination rate between parental locus j and k. The recombination frequency was estimated using the genetic map information as follows:

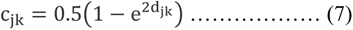

where d_jk_ is the map distance in morgan (M) between loci j and k (Haldane, 1919).

In addition, the UC was estimated as follows:

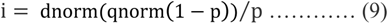

where μ_IS_ is the mean of the genetic value of the cross, i is the selection intensity, and 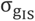 is the standard deviation estimated from Eq. 5. We calculated the standardized selection intensity using the following method in R environment:

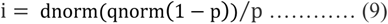

where p is the selected proportion.

To obtain the OHV of the parental combination, it was estimated as follows:

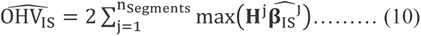

where n_Segments_ is the number of segments into which the genome is split, **H**^j^ is the matrix containing the four haplotype scores (0 or 1) of the two parental lines, 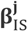 is the vector of marker effects of segment j estimated via Eq. 2 using training population. See Daetwyler et al. (2015) for details.

For all treatments, DH lines were made from the F_1_ to reduce computation time, and 15,000 individuals were generated for evaluation in the nursery. Visual selection with heritability of 0.1 was assumed across traits following (Gaynor et al. 2017). In the preliminary yield (PYT) trial, we evaluated 400 genotypes advanced from the nursery stage. These genotypes were assessed in two replicates across two locations. Based on the rank summation index, we identified superior genotypes and reintroduced them into the crossing block as parents. Cross combinations that generate progenies for the subsequent generations were determined using the different cross-selection metrics.

We further narrowed down the pool to 40 exceptional genotypes for the advanced yield trials. These 40 genotypes were evaluated in three replicates across seven locations and two years, providing us with comprehensive data on their performance for release as a variety.

Each treatment is independent, and the simulated breeding program spanned 40 years with a burn-in period of 10 years. Data regarding population mean, genetic gain, and genetic diversity were collected for the 10 to 40 years of the simulation, which was represented as 0 to 30 in the study. Each simulation treatment was replicated 50 times.

## Results

### Efficiency of multi-trait genomic inferred cross-selection metrics to simultaneously improve response to selection

When compared to other cross-selection metrics, the use of PMV as a cross-selection metric consistently results in a high genetic gain or response to selection for traits where an increase is expected, such as YLD and TKW (**Fig. 2** and **3**). Additionally, it effectively facilitates the desired selection response for other traits, such as optimal PH and DPM (**Fig. 4** and **5**). Except when using 30 parents, 200 crosses, and 75 progenies per cross, UC showed marginal gains over PMV in the medium term (15 to 20 years post burn-in) for YLD. A similar trend was observed when using the MeanPheno of non-related superior parents in a breeding scenario that involved 50 crosses derived from 50 parents, each with 300 progenies per cross, for both YLD (**Fig. 2**) and TKW (**Fig. 3**). In general, PMV outperforms other metrics across the different breeding strategies evaluated in this study.

**Figure 2:**
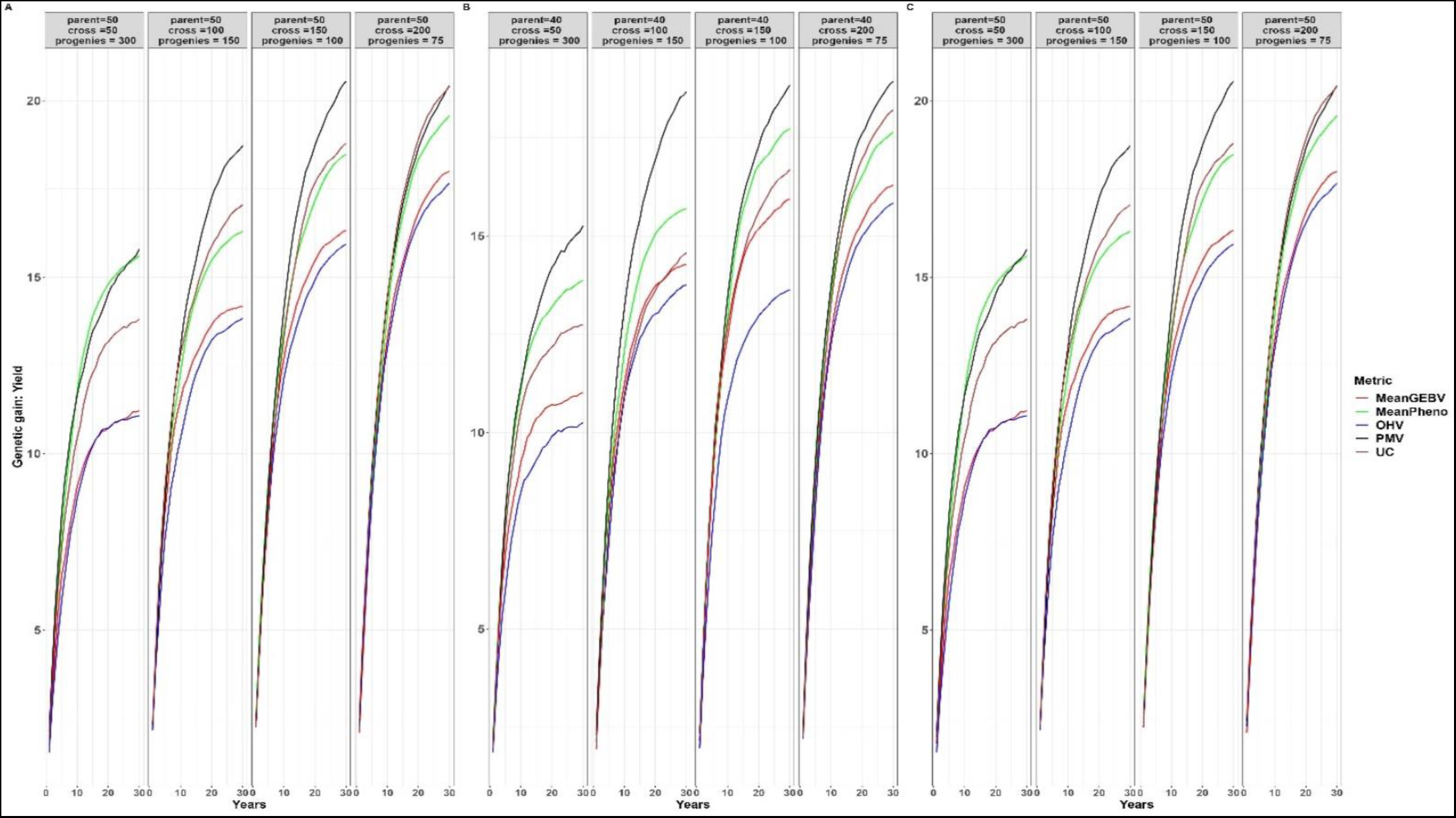
Genetic gains for different cross-selection metrics and different number of parents, crosses and progenies per cross for grain yield over 30 years post burn-in. The red line (MeanGEBV) highlights the genetic gain obtained using mean of the GEBV of the distantly related superior parents to select crosses, green line (MeanPheno) represents mean index selection value of the non-related superior parents, the blue line (OHV) is the optimal haploid value, black line (PMV) represents the posterior mean variance and brown line (UC) represent genetic gain observed using the usefulness criterion as cross-selection metric.

**Figure 3:**
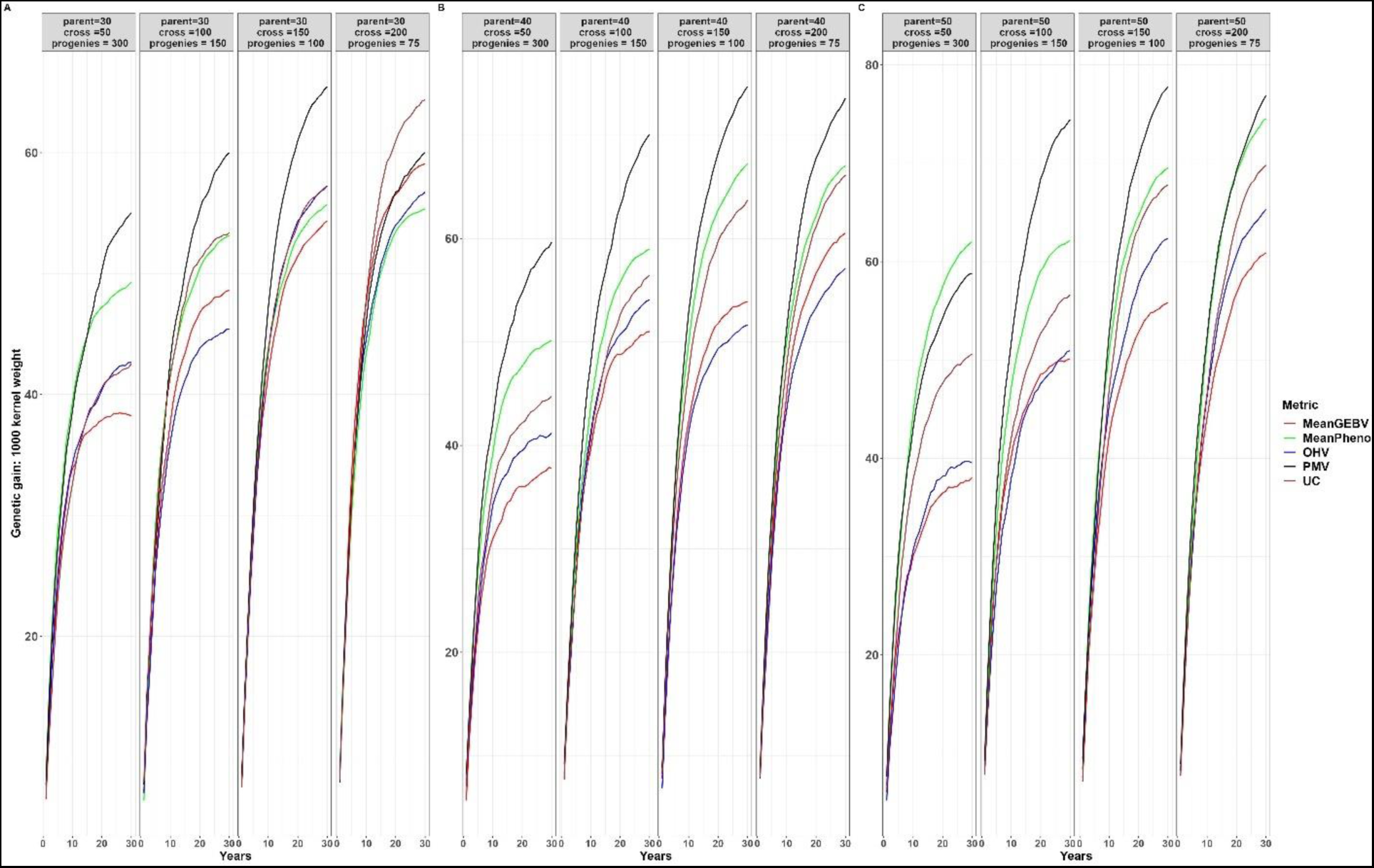
Genetic gains for different cross-selection metrics and different number of parents, crosses and progenies per cross for 1000 kernel weight over 30 years post burn-in. The red line (MeanGEBV) highlights the genetic gain obtained using mean of the GEBV of the distantly related superior parents to select crosses, green line (MeanPheno) represents mean index selection value of the non-related superior parents, the blue line (OHV) is the optimal haploid value, black line (PMV) represents the posterior mean variance and brown line (UC) correspond to the genetic gain observed using the usefulness criterion as cross-selection metric.

**Figure 4:**
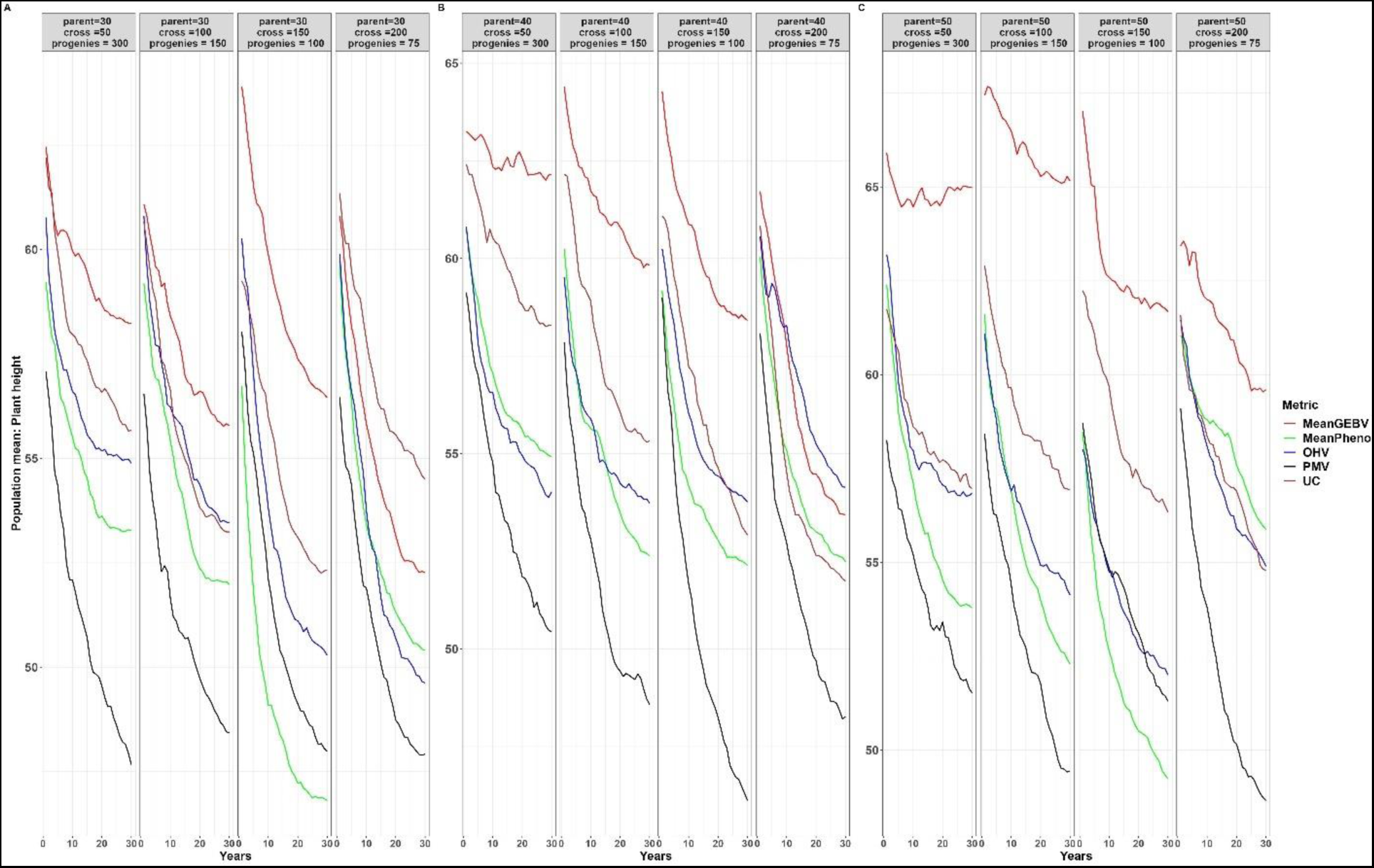
Population mean for different cross-selection metrics and different number of parents, crosses and progenies per cross for plant height over 30 years post burn-in. The red line (MeanGEBV) highlights the genetic gain obtained using mean of the GEBV of the distantly related superior parents to select crosses, green line (MeanPheno) represents mean of index selection value of the non-related superior parents, the blue line (OHV) is the optimal haploid value, black line (PMV) represents the posterior mean variance and brown line (UC) represents genetic gain observed using the usefulness criterion as cross-selection metric.

**Figure 5:**
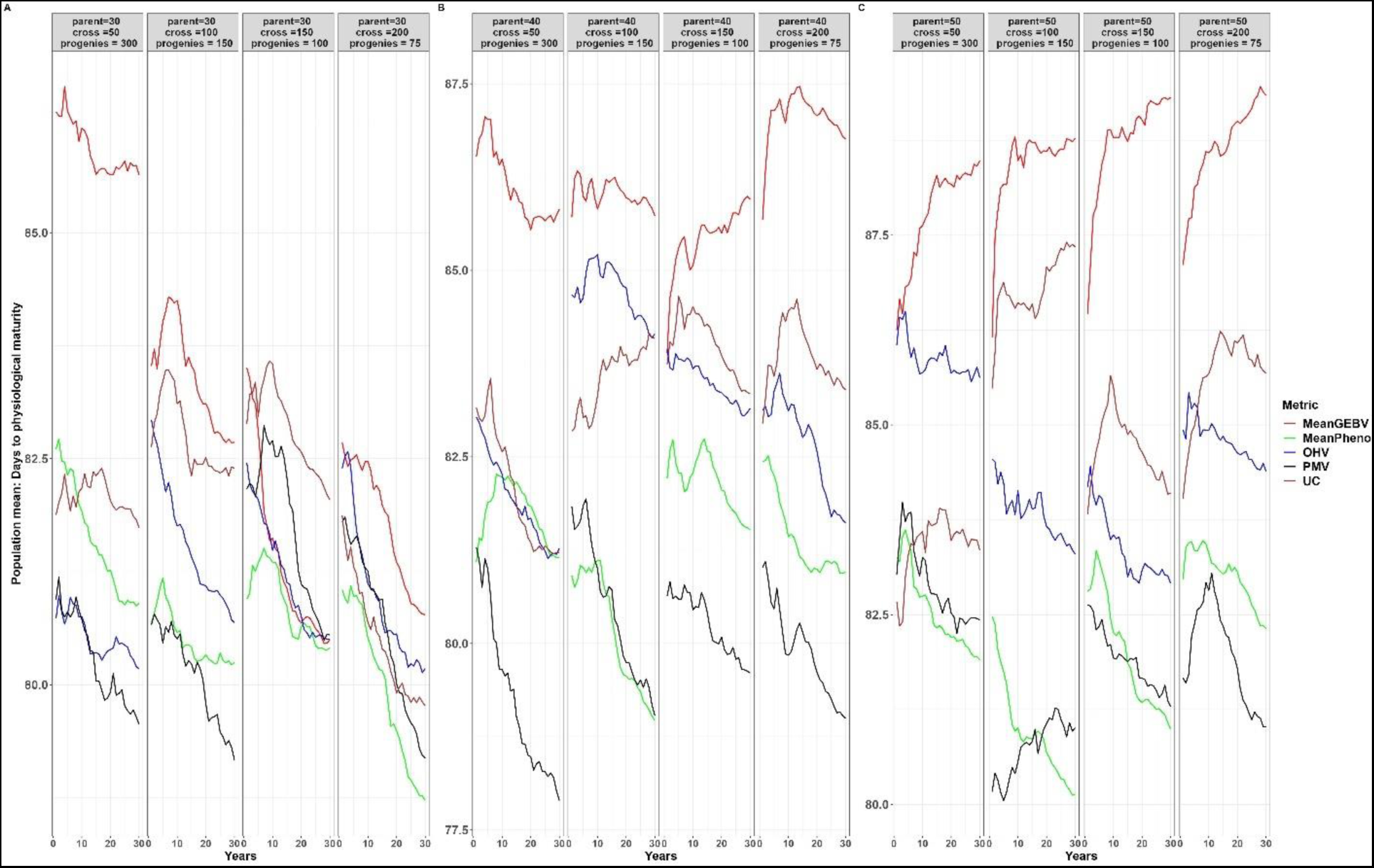
Population mean for different cross-selection metrics and different number of parents, crosses and progenies per cross for days to physiological maturity over 30 years post burn-in. The red line (MeanGEBV) highlights the genetic gain obtained using mean of the GEBV of the distantly related superior parents to select crosses, green line (MeanPheno) represents mean index selection value of the non-related superior parents, the blue line (OHV) is the optimal haploid value, black line (PMV) represents the posterior mean variance and brown line (UC) represents genetic gain observed using the usefulness criterion as cross-selection metric.

For example, employing PMV as the selection method, involving 40 parents, 50 crosses, and 300 progenies per parent, resulted in a higher genetic gain of 0.34% in the short term (1 to 10 years post-burn-in) and 8.56% in the long term (20 to 30 years post-burn-in) for YLD compared to using the MeanPheno of the parents as the selection metric **(Fig. 2)**. Using the same breeding method, the mean population for PH across the breeding cycles was 53.59cm for PMV and 56.65cm for MeanPheno **(Fig. 4)**. When compared to the founder population mean of 67.00cm, PMV efficiently select parents with optimal PH while sustaining gains for other primary traits. Moreover, the mean population for DPM was 79.25 days for PMV and 81.75 days for MeanPheno **(Fig. 5)**. In comparison to the founder population mean of 81 days, PMV led to a genetic gain of 2.19%.

Furthermore, the genetic gain for YLD improved by 7.53% in the short term and 16.32% in the long term when the number of crosses increased from 50 to 100 and the number of progenies per cross was reduced to 150 (**Fig. 2**). The mean population for PH using PMV was 51.52cm, compared to 54.89cm using MeanPheno (**Fig. 4**). There was no difference in the mean population of DPM, with PMV and MeanPheno having 80.44 and 80.16 days, respectively (**Fig. 5**). A similar trend was observed for other breeding scenarios. We emphasize the mean population values for both PH and DPM for simplicity, as these traits are expected to have optimal values in the long term, in contrast to the mean population values of the founder parents.

In all breeding scenarios and for every trait we considered, PMV has higher genetic variance when compared to all other selection methods, as depicted in **Fig. 6** and **Supp 2:4**. The magnitude of genetic diversity loss per unit of time was lower with PMV when compared to our baseline metric (MeanPheno), especially in the medium and long term. For instance, using the smallest number of parents (30) and crosses (50) in our study, we observed a substantial reduction in the magnitude of genetic variance after 30 years post-burn-in. When we use MeanPheno as our cross-selection metric, the genetic variance diminishes to 0.02, 0.38, 0.10, and 0.22 for the traits YLD, TKW, PH, and DPM, respectively. However, when we employ PMV, the genetic variance remains notably higher, at 0.091, 1.69, 0.59, and 0.99 for the same set of traits. In general, PMV consistently shows greater genetic diversity and slower loss of diversity over time.

**Figure 6:**
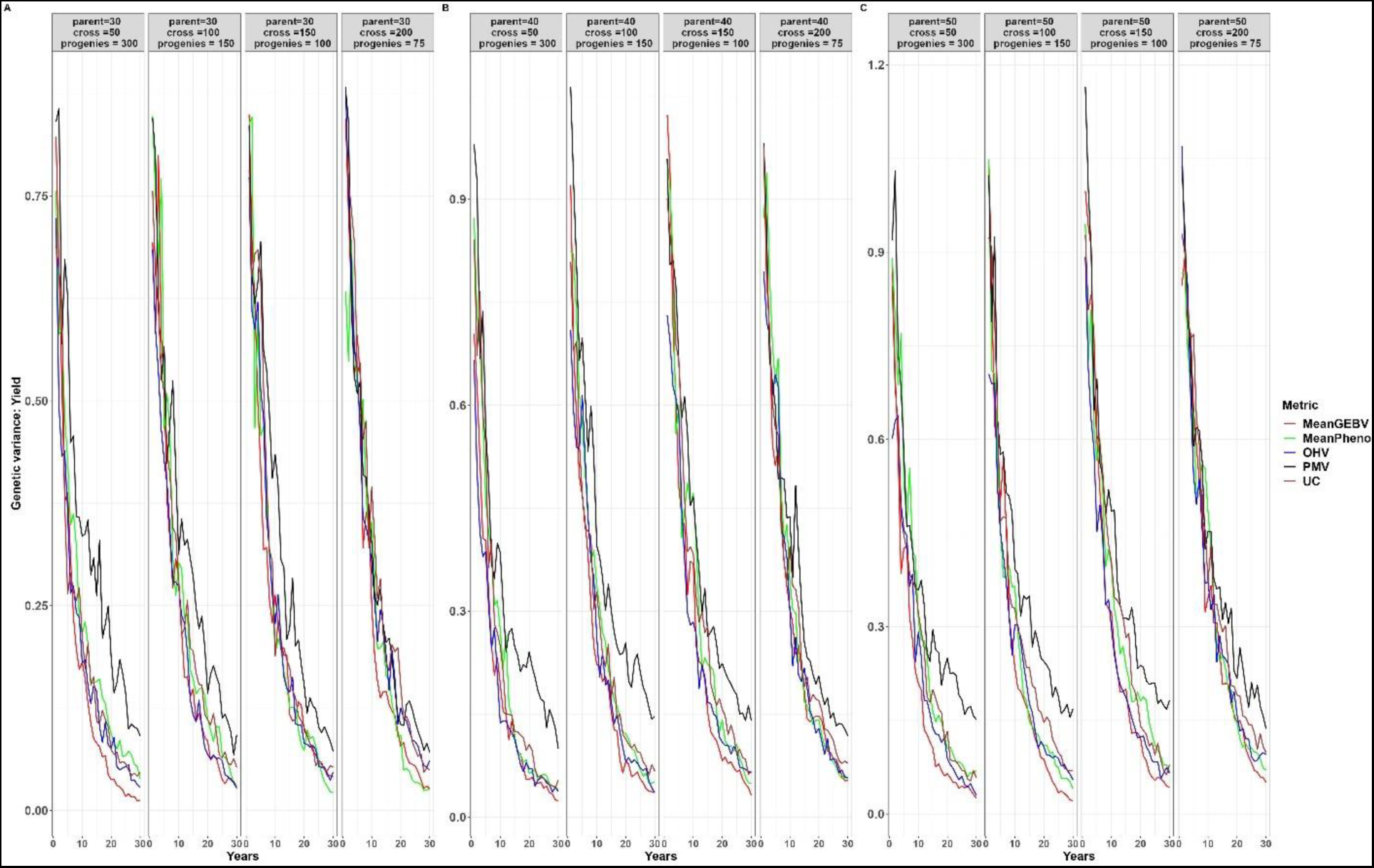
Genetic variance for different cross-selection metrics and different number of parents, crosses and progenies per cross for grain yield over 30 years post burn-in. The red line (MeanGEBV) highlights the genetic gain obtained using mean of the GEBV of the distantly related superior parents to select crosses, green line (MeanPheno) represents mean of index selection value of the non-related superior parents, the blue line (OHV) is the optimal haploid value, black line (PMV) represents the posterior mean variance and brown line (UC) represents genetic gain observed using the usefulness criterion as cross-selection metric.

### Number of Parents, Number of Crosses, and Number of progenies per cross

Genetic gain is influenced by a combination of factors (number of parents, crosses and progenies per cross) that appear to be interconnected, as depicted in Figure 7. We selected the PMV for assessing the number of parents, crosses, and population size due to its superior efficiency when compared to other metrics.

**Figure 7:**
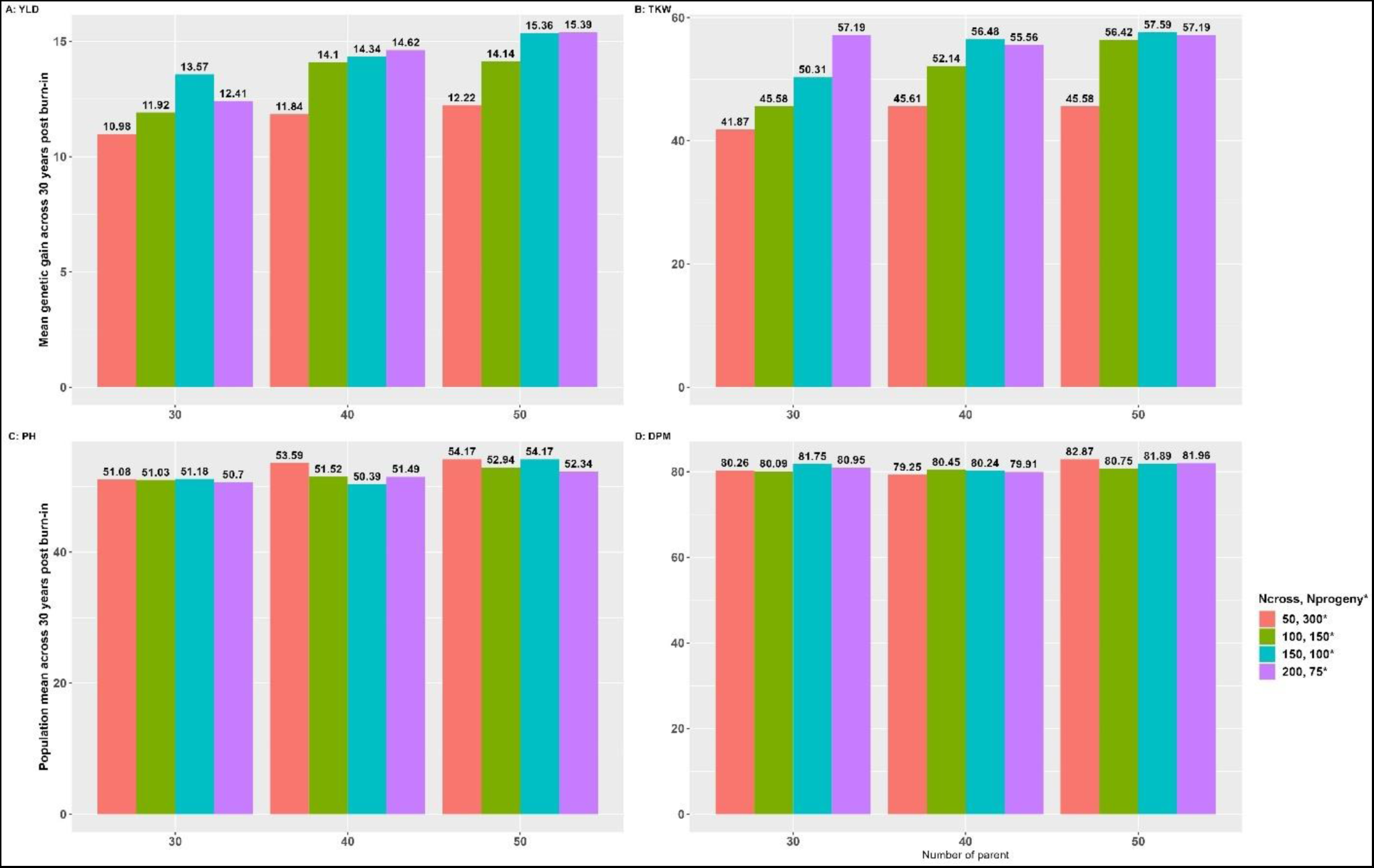
A and B represent genetic gains across 30 years post burn-in for grain yield (YLD) and 1000 kernel weight (TKW) and C and D represent the population mean across 30 years post burn-in for plant height (PH) and days to physiological maturity (DPM).

Increasing the number of crosses benefits from an increased number of parents but plateaus at 150 crosses (**Fig 7A:D**). Considering YLD, the genetic gain values were 10.98, 11.92, 13.57, and 12.41 for 50, 100, 150, and 200 number of crosses and 300, 150, 100, and 75 number of progenies per cross using 30 number of parents (**Fig. 7A**). The gain was only rapid when the number of crosses increased from 100 to 150. However, we observed diminishing returns when we further increased the number of crosses from 150 to 200. Similarly, when the number of parents was 40, the genetic gain increased from 11.84, 14.10, 14.34, and 14.62 for 50, 100, 150, and 200 crosses and 300, 150, 100, and 75 progenies per cross. Again, the gain from increasing the number of crosses beyond 150 was marginal. The same trend was observed with 50 parents, where the genetic gain increased from 12.22 for 50 crosses to 14.14 for 100 crosses, 15.36 for 150 crosses, and 15.39 for 200 crosses, while the number of progenies per cross remained at 300, 150, 100, and 75. Similar to the observation with 30 parents, there was no substantial improvement in genetic gain when increasing the number of crosses from 150 to 200.

Although there was a linear trend when the number of parents increased from 30 to 50, only marginal gains were observed when the number increased from 40 to 50. A similar trend was observed for TKW (**Fig. 7B**), except that the linear increase in the number of crosses consistently improved the genetic gain, especially for 30 parents.

Considering PH, we found that with 30 parents, there was no significant difference in the mean population (15.08, 15.03, 51.18, and 50.70cm) when increasing the number of crosses from 50 to 200 (**Fig. 7C**). However, it improved when compared to the mean population (67.00cm) of the founder population. When we increased the number of parents from 40 to 50, we observed a similar pattern, except for a few differences. In the case of DPM, increasing the number of parents and the number of crosses did not translate linearly to an improved response to selection (**Fig. 7D**).

## Discussion

One of the key drawbacks of genomic selection is the loss of genetic diversity when compared to traditional phenotypic selection in the long term (Jannink, 2010). In our study, we used stochastic simulation to evaluate the effectiveness of different genomic prediction cross-selection metrics to predict the usefulness or merit of a cross, particularly when aiming for simultaneous improvement across multiple traits. Furthermore, this study aimed to support the NDSU pulse breeding program in determining the optimal number of parents, crosses, and progenies per cross, taking into account the constraints posed by the current breeding budget and logistical considerations.

The observed selection gain for all traits in various breeding scenarios and the cross-selection metrics demonstrate the potential utility of using index selection as a derived phenotype within a genomic prediction framework. This approach can be especially valuable in situations where fitting multiple traits simultaneously might be computationally intensive or statistically challenging. In principle, this allows the selection of optimal parent combinations that strike a balance among multiple traits, taking into account their relative importance. Following a different rationale than the one described in our study, (Chung & Liao, 2022; Wolfe et al. 2021) also reported an increase in genetic gain for simultaneous improvement of multi-traits using genomic prediction to predict the merit of crosses. This approach is not without its limitations; its efficacy may vary with different index selection methods. The inability to identify optimal breeding parents that effectively balance multiple traits relative to their importance. Additionally, the numerical sensitivity of some selection index methods can lead to transformations or scaling of traits, potentially affecting their biological relevance.

Studies (Lehermeier et al. 2017; Allier et al. 2019; Wolfe et al. 2021) have reported that PMV serves as an unbiased predictor of progeny variance within bi-parental populations. This was attributed to PMV taking into account the haplotype of the parents, estimates of marker effects, and estimates of recombination frequencies between marker loci (Lehermeier et al., 2017; Allier et al., 2019). Souza & Sorrells (1991) suggested that the genetic gain achieved from a cross depends on the genetic variance of selected elite parents. Therefore, crosses with large genetic variance from the elite pool would theoretically generate a population with a favorable mean and contribute to increased genetic gain (Bernado, 2020, Moeinizade et al., 2019). This assumption will be invalid in an unstructured cross setting, where crossing parents is a mix of elite and poor performing lines. In practical breeding programs, the goal is often to maximize short-term gains while preserving long-term sustainability. Consequently, crosses between elite and poorly performing lines would be detrimental to achieving this objective (Cobb et al., 2019; Bernado, 2020). Our finding thus, suggest a path to balance short-term gain and long-term sustainability.

In a simulation study by (Zhong & Jannink, 2007), they observed predictive accuracy for progeny variance decreased as the number of QTL increased. In contrast, in our study, we did not observe a decrease in genetic gain for the traits we considered, despite variations in their genetic architecture. Furthermore, (Lehermeier et al. 2017) found no significant difference in the accuracy of progeny variance estimation when the number of QTL was 300 or fewer. This difference in outcomes could be attributed, at least in part, to the method we used to estimate marker effects, which was based on index selection rather than individual traits. This strategy also addressed the challenge highlighted by (Neyhart & Smith, 2019). They reported crosses with extreme population mean were accompanied by low genetic variance, while crosses with an intermediate population mean were associated with higher genetic variance. They explained that lines with similar genetic value will likely share alleles at the majority of quantitative trait loci (QTLs) underlying the trait; which accounts for the observed variation. However, this was not a concern in our proposed strategy because we are interested in predicting the variance of the index selection rather than individual traits, thus eliminating the chance of crossing poor lines with elite lines. For example, considering the smallest number of parents (30) and crosses (50) in our study, along with the increased genetic gain, PMV showed 4.56, 4.44, 5.90, and 4.50 times greater genetic diversity and a slower rate of genetic diversity for the traits YLD, TKW, PH, and DPM compared to the base metric (MeanPheno; average index value of the superior non-related parents) after 30 years post burn-in.

The inconsistent genetic gain observed when the UC was used for selection decision might be due to the dependency of the UC on selection intensity and trait heritability. Unsurprisingly, Lehermeier et al. (2017) found that selection of crosses based on UC is more advantageous with increased selection intensity and high heritability. Unexpectedly, OHV performed less favorably, our results was consistent with previous studies (Han et al., 2017; Lehermeier et al., 2017). Theoretically, OHV assumes an infinite number of progenies per cross and selection intensity (Daetwyler et al., 2015; Lehermeier et al. 2017), an assumption not met in our study. Furthermore, we did not fine-tune the number of segments where the absence of recombination is assumed, which is crucial for arriving at an optimal value.

Our results showed that the number of parents involved in crossing has a significant impact on both the population mean and genetic variance across the breeding cycles. In particular, when fewer parents are involved, we observed a slower rate of genetic improvement and an elevated risk of losing genetic diversity. This is primarily attributed to the lack of unique crosses, especially with the increased number of crosses. Additionally, alleles that are lost as a result of limited parental diversity was not regained in subsequent generations. This leads to the observed rapid decline in genetic diversity, particularly in the long term, resulting in reduced genetic gains compared to scenarios where a greater number of parents are involved.

Recently, Sabadin et al. (2022) also emphasized the relationship between the number of parents and the effective population size (*N_e_*). In the simulation study, the author reported greater resilience to the loss of genetic variance over the long term, involving 48 parents compared to 24 parents. The decrease in genetic gain when few individuals are used to form the next generation suggests that the effect of genetic drift may far outweigh the effect of response to selection (Bernardo, 2020). Therefore, selecting the appropriate number of parents for a breeding program is a pivotal factor in either accelerating or decelerating genetic progress, which directly impacts the program overall success (Cobb et al., 2019; Covarrubias-Pazaran et al., 2021).

Considering the constraints on the breeding program, such as limitations on the number of lines that can be evaluated, it becomes crucial to identify the balance between maximizing genetic gain and preserving valuable genetic diversity. Generally, increasing the number of crosses enhances genetic gain and reduces the risk of genetic drift; however, there is not much gain to achieve much larger than increasing the crosses from 50 to 150 with a population size of 300 to 100 at any given number of parents. Similarly, Covarrubias-Pazaran et al. (2021) also reported a sustained genetic gain in the long term with an increased number of crosses and fewer progenies per cross, but with diminishing returns to additional crosses with fewer parents. Therefore, to achieve sustainable genetic progress in a breeding program, especially for small breeding programs, caution should be exercised when determining the optimal number of parents, crosses, and progenies per cross.

## Conclusion

We presented a simple but efficient approach to identifying crosses that simultaneously improve the genetic gain of multi-trait using index selection of the parents, parental haplotypes, marker effects, and recombination frequencies between marker loci. We proposed the use of this cross-selection strategy in a breeding program implementing genomic selection to guarantee sustainable genetic improvement. For continuous delivery of varieties to the market and population improvement, the use of genetic simulation to guide optimal resource allocations and the design of crossing blocks is highly recommended. This study is not without its limitations. The underlying assumptions and simulated genetic parameters were tailored to the NDSU pulse breeding program, which might limit its generalization to other programs. To validate these results and extend it relevance to diverse breeding programs, empirical data should be used in multiple breeding programs. Nevertheless, our results serve as a guide for continuous genetic improvement in any breeding program.

## Data availability statement

No empirical data generated in this study and the procedure to replicate this study has been well documented. Further inquiries can be directed to the corresponding authors.

## Author Contributions

SA conceptualized the study, wrote the code, performed the analyses, and wrote the manuscript. NB conceptualized the study and contributed to the writing of the manuscript. All authors edited, reviewed, and approved the manuscript.

## Conflict of Interest

The authors declare that the study was conducted in the absence of any commercial or financial relationships that could be construed as a potential conflict of interest.

## Acknowledgments

This research was supported through funding from USDA-NIFA (Hatch Project ND01513) and the North Dakota Department of Agriculture through the Specialty Crop Block Grant Program (19–429).

---

**Supplementary 1:**
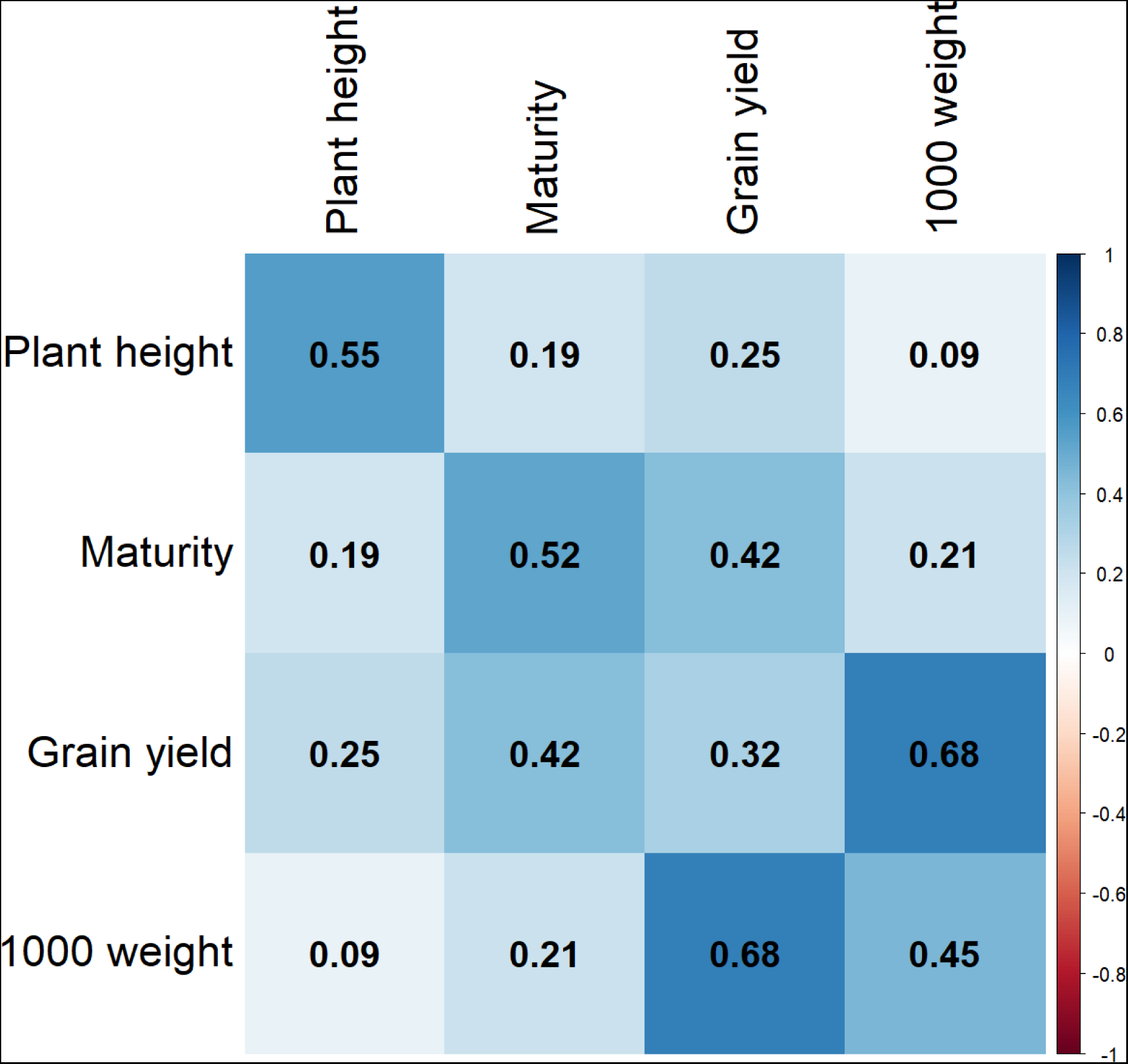
Genetic correlation of grain yield, 1000 kernel weight, plant height and days to physiological maturity. The diagonal represents heritability for each trait.

**Supplementary 2:**
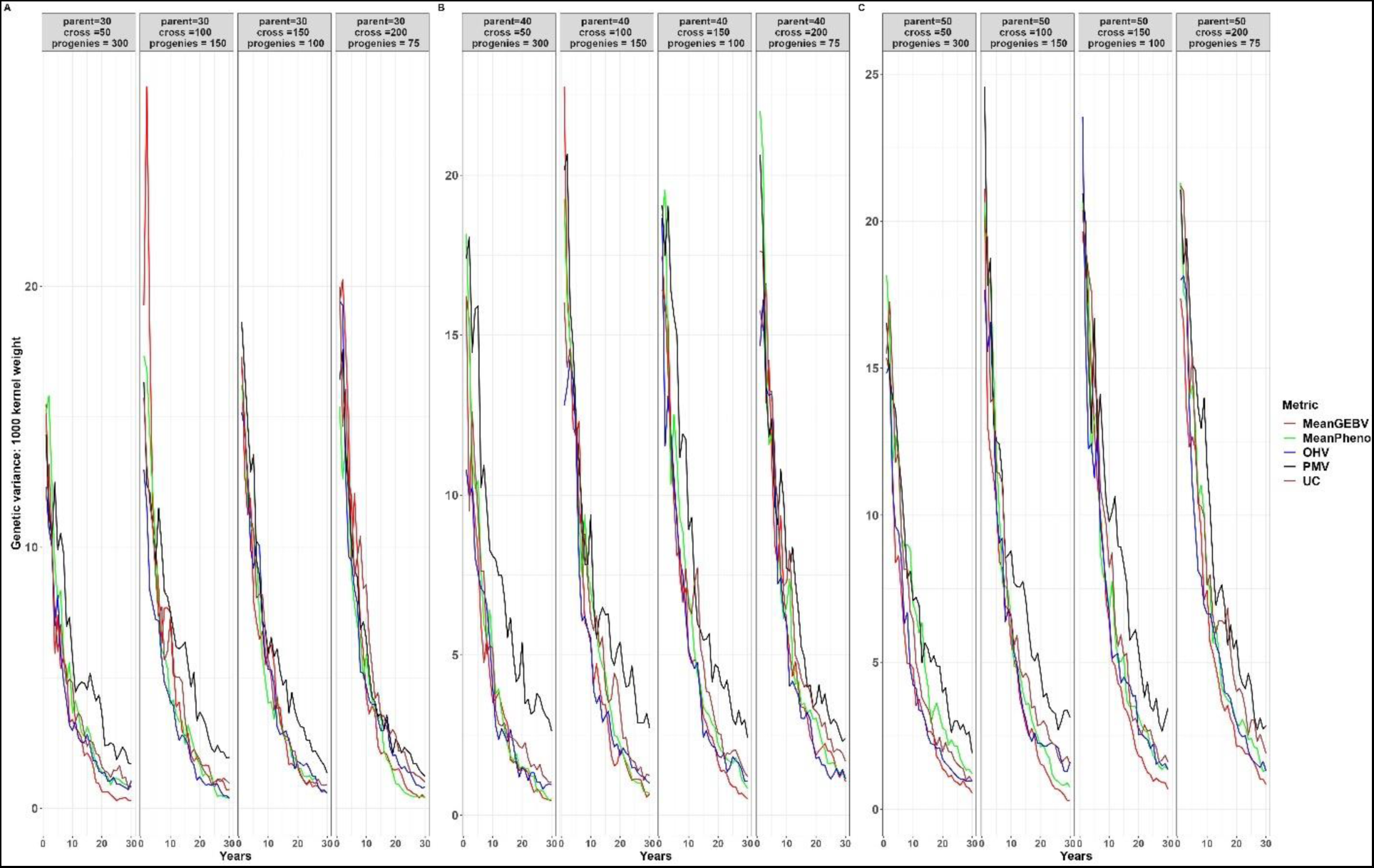
Genetic variance for different cross-selection metrics and different number of parents, crosses and progenies per cross for 1000 kernel weight over 30 years post burn-in. The red line (MeanGEBV) highlights the genetic gain obtained using mean of the GEBV of the distantly related superior parents to select crosses, green line (MeanPheno) represents mean index selection value of the non-related superior parents, the blue line (OHV) is the optimal haploid value, black line (PMV) represents the posterior mean variance and brown line (UC) represent genetic gain observed using the usefulness criterion as cross-selection metric.

**Supplementary 3:**
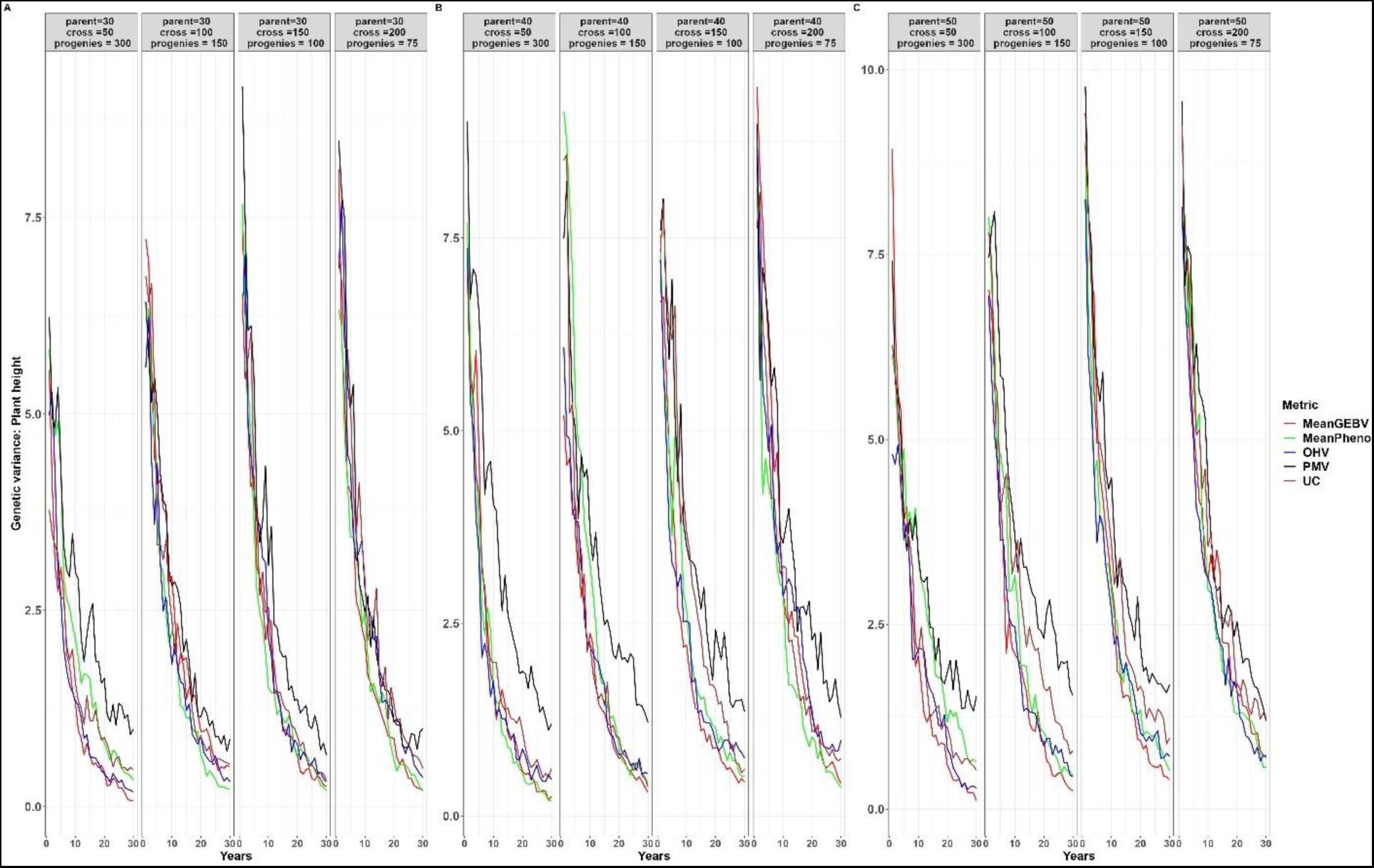
Genetic variance for different cross-selection metrics and different number of parents, crosses and progenies per cross for plant height over 30 years post burn-in. The red line (MeanGEBV) highlights the genetic gain obtained using mean of the GEBV of the distantly related superior parents to select crosses, green line (MeanPheno) represents mean index selection value of the non-related superior parents, the blue line (OHV) is the optimal haploid value, black line (PMV) represents the posterior mean variance and brown line (UC) represent genetic gain observed using the usefulness criterion as cross-selection metric.

**Supplementary 4:**
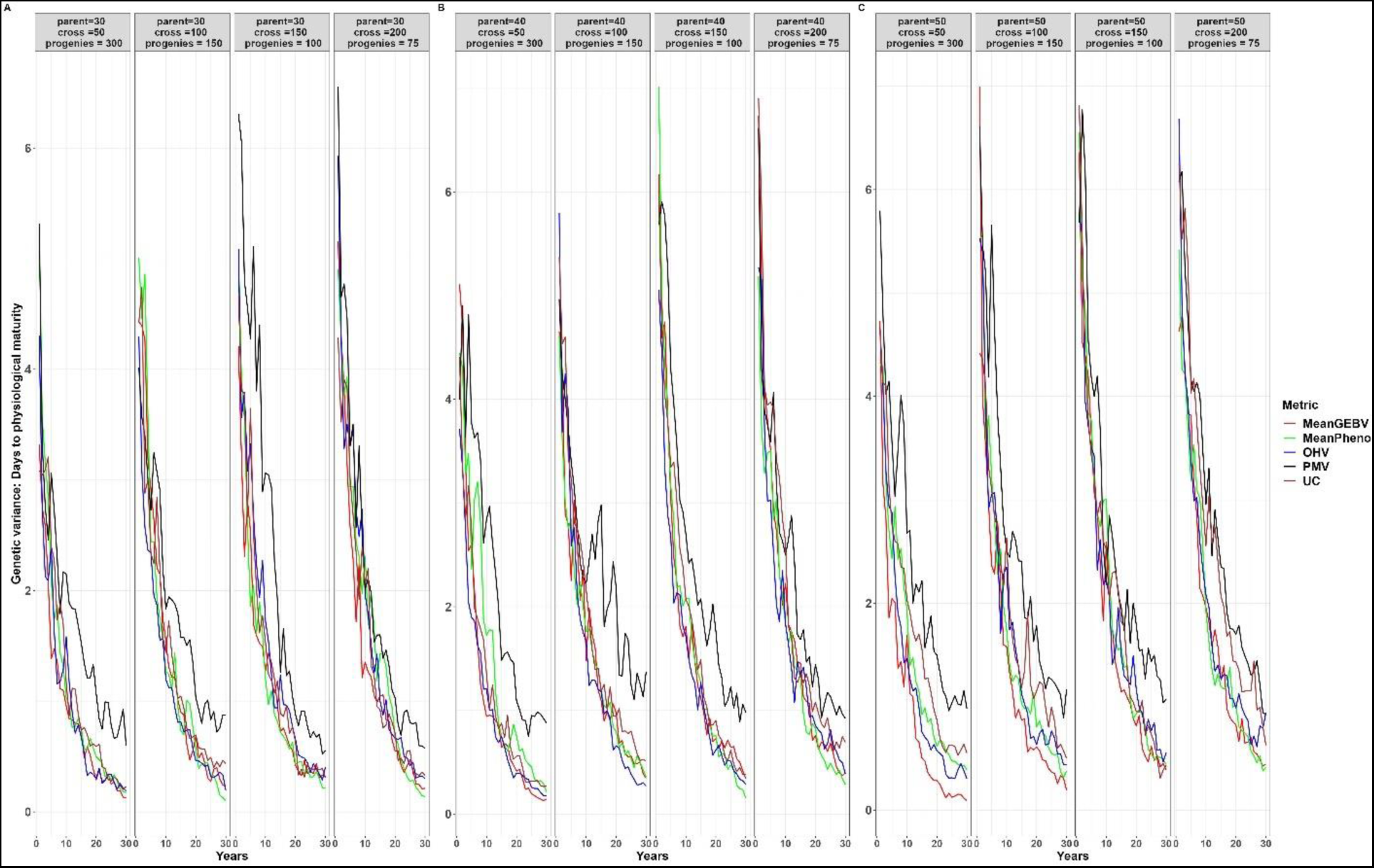
Genetic variance for different cross-selection metrics and different number of parents, crosses and progenies per cross for days to physiological maturity over 30 years post burn-in. The red line (MeanGEBV) highlights the genetic gain obtained using mean of the GEBV of the distantly related superior parents to select crosses, green line (MeanPheno) represents mean index selection value of the non-related superior parents, the blue line (OHV) is the optimal haploid value, black line (PMV) represents the posterior mean variance and brown line (UC) represents genetic gain observed using the usefulness criterion as cross-selection metric.

## Notes

### Competing Interest Statement

The authors have declared no competing interest.

### Summary of Updates

To update Equation 5 in the formal document.

